# The transcription factor E2A drives neural differentiation in pluripotent cells

**DOI:** 10.1101/736033

**Authors:** Chandrika Rao, Mattias Malaguti, John O. Mason, Sally Lowell

## Abstract

The intrinsic mechanisms that link extracellular signalling to the onset of neural differentiation are not well understood. In pluripotent mouse cells, BMP blocks entry into the neural lineage via transcriptional upregulation of Inhibitor of Differentiation (Id) factors. We have previously identified that the major binding partner of Id proteins in pluripotent cells is the basic helix-loop-helix (bHLH) transcription factor (TF), E2A. Id1 can prevent E2A from forming heterodimers with bHLH TFs or from forming homodimers. Here, we show that overexpression of a forced E2A homodimer is sufficient to drive robust neural commitment in pluripotent cells, even under non-permissive conditions. Conversely, we find that E2A null cells display a defect in their neural differentiation capacity. E2A acts as an upstream activator of neural lineage genes, including *Sox1* and *Foxd4*, and as a repressor of Nodal signalling. Our results suggest a crucial role for E2A in establishing neural lineage commitment in pluripotent cells.

## Introduction

Following establishment of the primary germ layers during gastrulation, the embryonic neural plate is specified from the anterior ectoderm at approximately embryonic (E) day 7.5 in a process known as neural induction (Tam and Zhou, 1996). Pluripotent embryonic stem (ES) cells recapitulate central features of this process when differentiated in culture (Ying et al., 2003b), providing a useful system in which to study the mechanisms guiding these initial cell fate decisions during development. Whilst the key extracellular signalling pathways that inhibit neural lineage commitment have long been established, less progress has been made in identifying the downstream effectors of these pathways. In particular, inhibition of BMP signalling is crucial for establishment of the neuroectoderm (Di-Gregorio et al., 2007; Harland, 2000). The finding that BMP signalling inhibits neural differentiation via transcriptional upregulation of Id1 (Ying et al., 2003a) provides compelling evidence that Id proteins block the activity of a factor which would otherwise trigger the onset of neural commitment.

Id proteins lack a DNA binding domain and function primarily as dominant negative inhibitors of bHLH transcription factors, binding to and preventing them from forming functional dimers (Norton, 2000). It has previously been reported that Id1 indirectly inhibits the activity of the bHLH transcription factor, *Tcf15*. However, *Tcf15* appears to play a role in general priming for differentiation by enabling morphological changes, and does not have a specific instructive role in neural commitment (Davies et al., 2013) (Lin and Tatar, in prep). We have also found that E-cadherin acts downstream of BMP to help suppress neural commitment (Malaguti et al., 2013), but it is not known if or how Id1 is mechanistically linked with this process. Id1 is also reported to block the activity of the epigenetic regulator, Zrf1, preventing derepression of neural genes in ES cells (Aloia et al., 2015). Zrf1 overexpression alone, however, is not sufficient to drive expression of these genes, suggesting a requirement for additional factors to initiate neural differentiation in ES cells.

We have previously identified the E2A gene products, E47 and E12, as the main binding partners of Id1 in ES cells (Davies et al., 2013). E2A (also known as *Tcf3 -* not to be confused with Tcf7L1, which is also commonly known as Tcf3) belongs to the E-protein family of bHLH transcription factors, which also includes HEB (*Tcf12*) and E2-2 (*Tcf4*). E2A is able to regulate transcription of its target genes either by homodimerisation, or by heterodimerisation with class II bHLH transcription factors, such as the proneural factors Ascl1 and Neurogenin1/2 (Murre et al., 1989). Whilst E2A-bHLH heterodimers are well-established regulators of several fate determination processes, including neuronal subtype specification (Imayoshi and Kageyama, 2014), E2A homodimers have only been identified to function in the context of B-cell development (Shen and Kadesch, 1995), and it is not currently known whether this homodimer could also operate to control cell fate in other contexts.

E2A knockout mouse models have thus far failed to identify any overt gastrulation defects, with a failure of B-cell specification being the only major phenotype described to date (Bain et al., 1994; Zhuang et al., 1994). More recent analysis of these models, however, has noted that knockout mice have a significantly reduced brain size compared to their wild-type counterparts (Ravanpay and Olson, 2008), suggesting that a more in-depth investigation into the role of E2A during the earlier stages of development may be required to uncover subtle neural differentiation defects. In *Xenopus* embryos, loss of E2A has been associated with inhibition of gastrulation (Yoon et al., 2011). Additionally, E2A and HEB have been shown to be cofactors of the Nodal signalling pathway, both in human ES cells and in *Xenopus* (Yoon et al., 2011), with E2A playing a dual role to directly repress the Nodal target gene, *lefty*, during mesendoderm specification in *Xenopus*, whilst also driving the expression of dorsal cell fate genes (Wills and Baker, 2015).

In this study, we investigate a potential role for E2A homodimers in neural fate commitment. We first set out to characterise the expression of E2A during early neural differentiation by generation of an endogenously tagged E2A-V5 ES cell line. Using a gain-of-function approach we find that overexpression of a forced E2A homodimer, but not monomer, is sufficient to drive neural commitment of ES cells, even in the presence of serum, providing novel mechanistic insight into the molecular events unfolding downstream of Id1 during neural commitment. CRISPR/Cas9 targeting of E2A and HEB loci to generate single and double E-protein knockout ES cell lines additionally reveals that E-protein deficiency compromises the neural differentiation ability of ES cells. RNA-sequencing analysis further confirmed that E2A is positioned upstream of the expression of several neural lineage-associated genes, including *Sox1* and *Foxd4*, and may additionally play a role in suppressing Nodal signalling during neural differentiation. We therefore propose that the E2A homodimer is a key intrinsic regulator of neural fate commitment in pluripotent ES cells.

## Results

### E2A is expressed heterogeneously in pluripotent ES cells and throughout neural differentiation

To characterise the expression of E2A during early neural differentiation we first examined the temporal expression of *E2A* mRNA in ES cells plated under standard neural monolayer conditions (Ying et al., 2003b) by qRT-PCR. *Sox1* is the earliest specific marker of neuroectoderm in mice (Pevny et al., 1998) and is therefore used to follow neural fate acquisition in ES cells. In line with previously published data (Aiba et al., 2009; Ying et al., 2003b), we observed that expression of the negative regulator of E2A, *Id1*, is rapidly downregulated at the onset of differentiation (***Figure 1A***). *E2A* expression, however, remains fairly constant during this initial period. As E2A is regulated by Id1 on the protein level, rather than the transcriptional level, we generated an endogenously tagged ES cell line (***Figure 1B***) using CRISPR/Cas9 targeting to follow the expression of E2A protein during differentiation. Based on a strategy previously developed to tag neural stem cells with high efficiency (Dewari et al., 2018), guide RNA (sgRNA) was designed to cut proximal to the stop codon in the 3’UTR of *E2A (****Supplementary Figure 1A)***, and was co-transfected into wild-type ES cells with recombinant Cas9 protein (rCas9) and a single-stranded donor DNA template (ssODN) encoding the V5 tag flanked by homology arms (***Supplementary Figure 1B***). Clonal lines were isolated from the bulk population, and individual clones were subsequently screened by PCR genotyping (***Supplementary Figure 1C***) and analysed by immunostaining (***Figure 1C***) for the V5 fusion protein. Sanger sequencing confirmed error-free insertion of V5 into the C-terminus of the *E2A* locus by homology-directed repair (HDR) (***Supplementary Figure 1D***).

**Figure 1.**
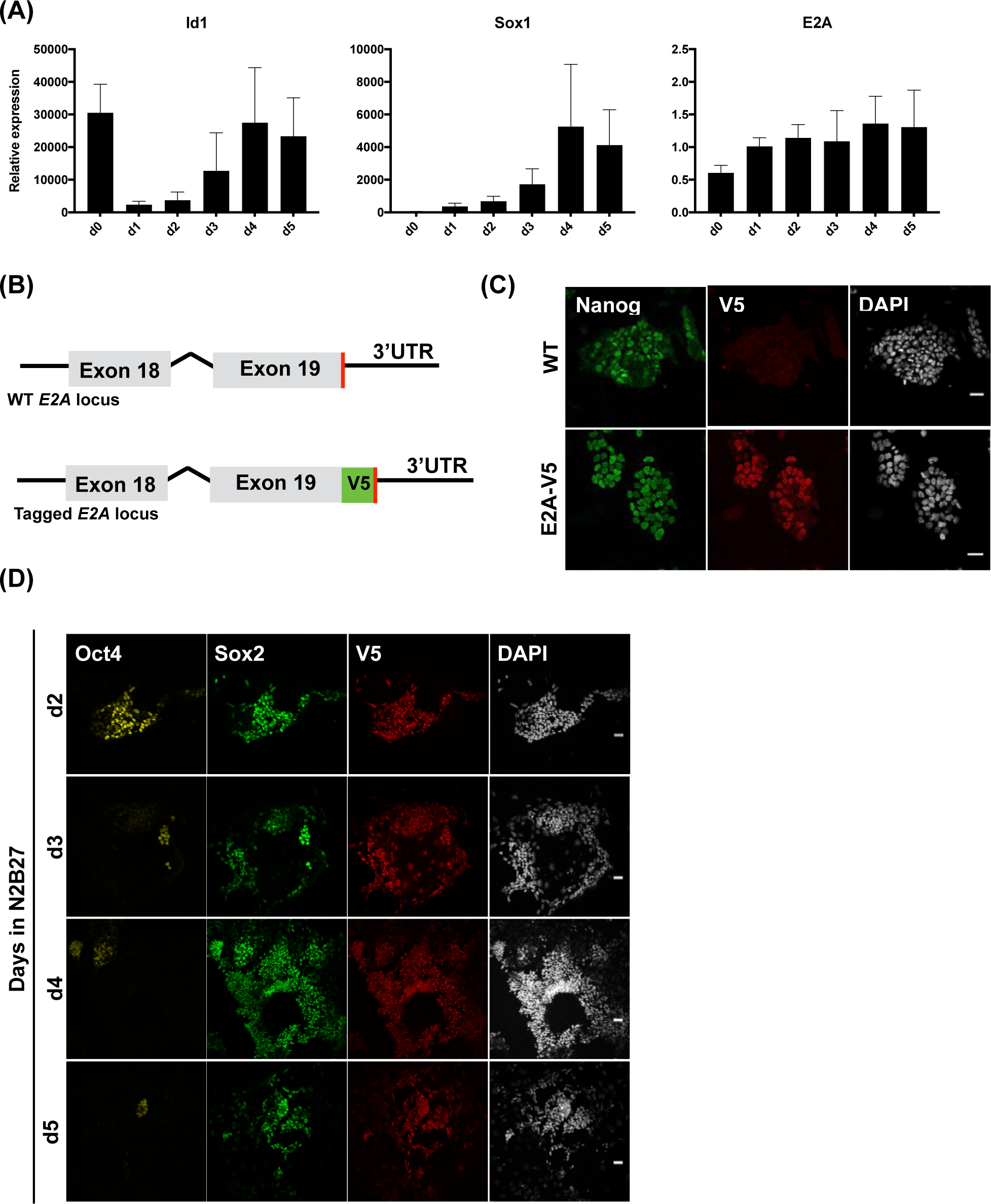
Endogenous tagging reveals E2A is expressed heterogeneously in pluripotent cells. **(A)** qRT-PCR analysis of pluripotent ES cells plated under standard neural differentiation conditions. Error bars represent standard deviation of the mean of 3 biological replicates. **(B)** Schematic of WT and tagged *E2A* locus. The V5 epitope tag was knocked in to the 3’UTR of the endogenous *E2A* locus. V5 tag is shown in green and stop codon in red. **(C)** Immunostaining of parental wild-type ES cells and epitope-tagged E2A-V5 ES cells. Cells co-stained for V5 tag, the pluripotency marker Nanog, and nuclear marker DAPI. **(D)** Immunostaining of E2A-V5 cells during days 2-5 neural differentiation. Cells stained for Oct4 and Sox2 to enable identification of cells committing to the neural lineage (Oct4^−^/Sox2^+^). Scale bar: 30 µm

Immunostaining of the resulting E2A-V5 cell line using an antibody against the V5 tag revealed that E2A expression is heterogeneous in pluripotent ES cells in LIF/serum culture (***Figure 1C***). When E2A-V5 ES cells are plated under neural differentiation conditions, E2A expression remains high and retains a heterogeneous expression pattern as cells exit the pluripotent state (Oct4^+^Sox2^−^) and progressively commit to the neural lineage (Oct4^−^Sox2^+^) (***Figure 1D***).

### E2A homodimers promote neural fate commitment under self-renewal conditions

Having observed that E2A expression is dynamic during the early stages of differentiation we next wanted to address whether E2A could play an instructive role during this cell fate transition. To determine whether E2A is sufficient to promote neural differentiation we generated doxycycline-inducible ES cell lines to overexpress either monomeric E2A, or a forced E2A homodimer in which two E2A sequences are tethered by a flexible amino acid linker (***Figure 2A***). This forced dimer strategy not only renders E2A more resistant to inhibition by Id, but also favours the formation of E2A homodimers due to the physical proximity of the two molecules (Neuhold and Wold, 1993). FLAG-tagged E47 monomer or forced homodimer constructs were placed under the control of a tetracycline-response element and introduced into ES cells containing an inducible cassette exchange locus upstream of the HPRT gene (Iacovino et al., 2013). When expression of the two E2A constructs was induced by doxycycline (dox) in LIF/serum culture - conditions which are inhibitory for neural differentiation - we found that overexpression of the forced E2A homodimer, but not the monomer, elicited a robust upregulation of Sox1 (***Figure 2B,D and Supplementary Figure 2***). Coincident with the peak of *Sox1* expression on day 2, we also observed that cells overexpressing the homodimer lose the domed colony morphology typical of ES cells, and instead spread out to cover the dish. We evaluated that approximately one-third (34%) of cells were Sox1^+^Oct4^−^ after four days of culture (***Figure 2C***), indicative of neural commitment. qRT-PCR analysis revealed that forced expression of E2A homodimers also enhanced expression of the neural marker *N-cadherin* (*Cdh2*), downregulated *E-cadherin* (*Cdh1*) expression, and caused a transient upregulation of the epiblast marker *Fgf5 (****Figure 2D***). Overexpression of both the monomeric and forced homodimer forms of E2A also appeared to stimulate a feedback response, causing upregulation of the negative regulator of E2A, *Id1*, as has been observed previously (Bhattacharya and Baker, 2011).

**Figure 2.**
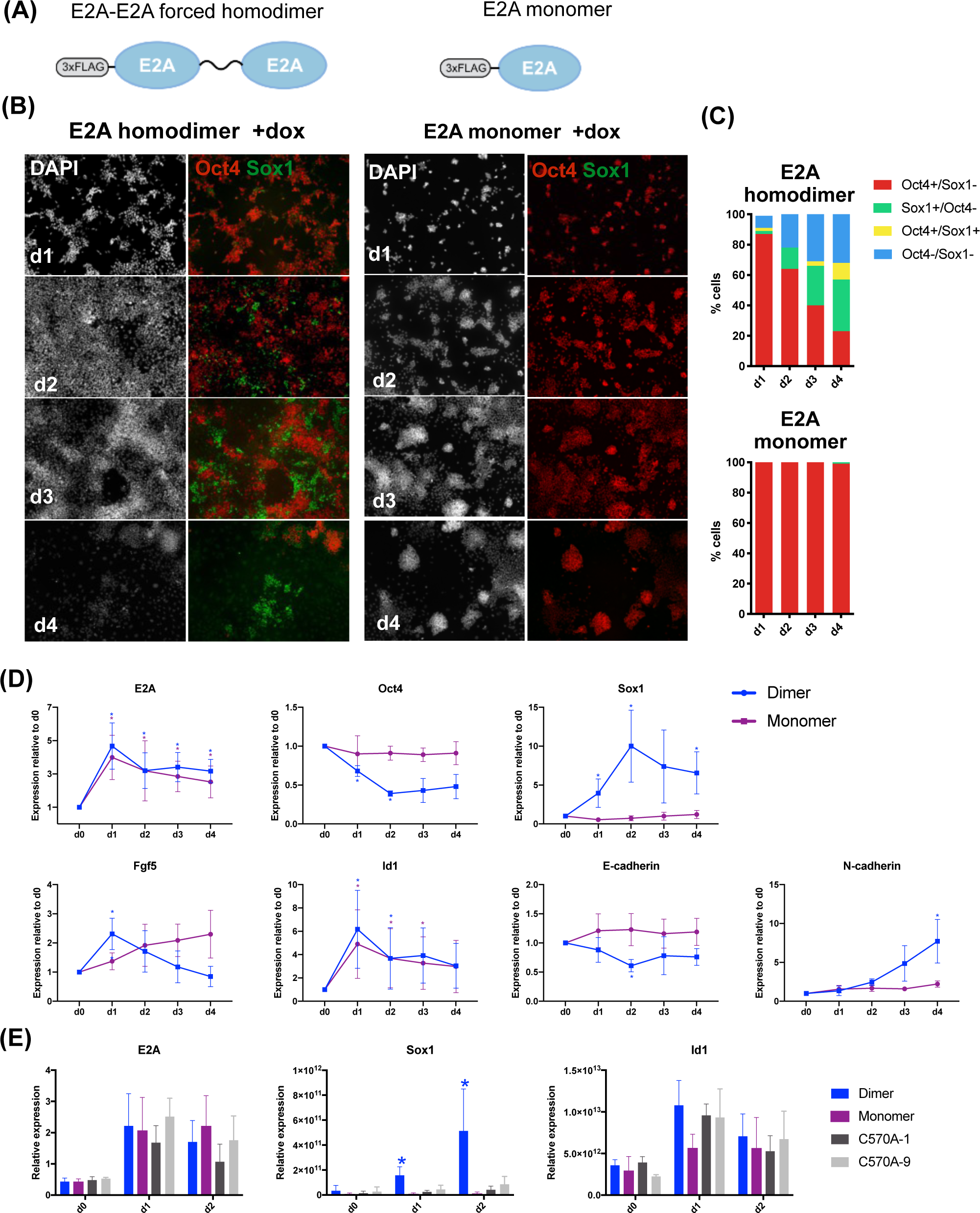
An E2A forced homodimer drives Sox1 expression under self-renewal conditions. **(A)** Schematic representation of monomeric E2A and forced E2A homodimer constructs. **(B)** Immunostaining of inducible E2A forced homodimers and monomers cultured in LIF/ serum+dox. **(C)** Quantification of immunostaining for E2A monomers and homodimers cultured in LIF/serum +dox. 500-5000 cells were manually scored for each timepoint. **(D)** qRT-PCR analysis of dox-inducible overexpression of E2A monomers and homodimers in LIF/serum. Expression is normalised to day 0 minus dox controls. **(E)** qRT-PCR analysis of two mutant forced homodimer clones (C570A-1 and C570A-9) induced in LIF/serum +dox. Error bars represent standard deviation of the mean of a minimum of 3 biological replicates. *p<0.05 by Student’s t-test.

It has been reported that E2A homodimers are stabilised by an intermolecular disulphide bond formed between cysteine residues located in helix one of the bHLH domain (Benezra, 1994) (***Supplementary Figure 3A***). To confirm that it was specifically E2A homodimers, and not E2A-bHLH heterodimers, that are driving neural commitment in ES cells, we generated E2A forced dimer cell lines in which cysteine-570 is mutated to alanine in both E2A sequences (***Supplementary Figure 3B***), aiming to disrupt the formation of this covalent bond and therefore destabilise homodimer formation. Two such mutant lines, C570A-1 and C570A-9, were stimulated with doxycycline in LIF/serum for two days. qRT-PCR analysis of both mutant and control inducible lines showed that whilst C570A mutants were still able to upregulate *Id1*, they were no longer able to upregulate expression of *Sox1* (***Figure 2E***), suggesting that this single amino acid substitution did, indeed, ablate the ability of E2A to drive differentiation. Taken together these findings demonstrate that overexpression of E2A homodimers is sufficient to override the potent inhibitory effect of serum in non-permissive culture conditions, consistent with a role for E2A homodimers in initiating neural fate commitment.

### E2A^−/−^ and E2A^−/−^HEB^−/−^ knockout ES cells display compromised neural lineage commitment ability

We next set out to evaluate whether E2A is necessary for neural differentiation. To generate E2A knockout ES cells, we used CRISPR/Cas9 to target exon 3 of the *E2A* gene locus, disrupting both *E12* and *E47* transcripts. Due to the high degree of functional compensation observed between E-protein family members (Zhuang et al., 1998), which could mask potential differentiation phenotypes, we also generated E2A^−/−^HEB^−/−^ double E-protein knockout ES cell lines by using a similar strategy to target exon 9 of *HEB* in the E2A^−/−^ cell line (***Supplementary Figure 4A***). Targeted lines were validated by western blot analysis and sequencing of the targeted loci (***Supplementary Figure 4B-F***).

To first determine whether E-protein knockout cells retain characteristic features of pluripotent stem cells, we assessed cell morphology and gene expression in LIF/serum conditions. Colony morphology was indistinguishable between the parental E2A-V5 cell line and E2A^−/−^ and E2A^−/−^HEB^−/−^ cells, and immunostaining showed similar levels of expression of the pluripotency markers Oct4 and Nanog between knockout and parental cells (***Figure 3A***), indicating normal self-renewal. qRT-PCR analysis of pluripotency and lineage markers in LIF/serum further confirmed that there were no differences in expression levels between parental and knockout cell lines (***Figure 3B***).

**Figure 3.**
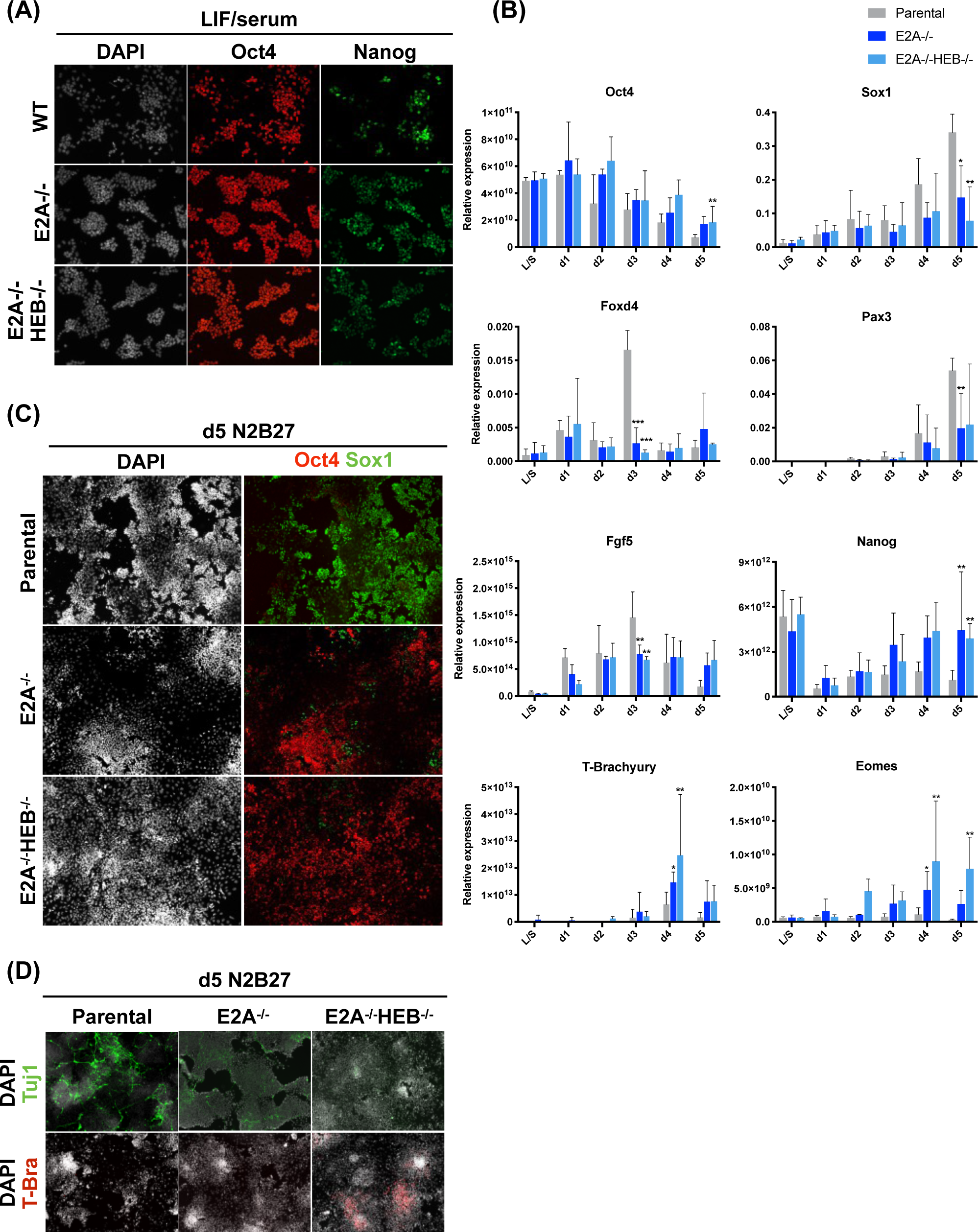
E2A^−/−^ and E2A^−/−^HEB^−/−^ ES self-renew normally but display defects in early neural commitment. **(A)** Immunostaining of WT, E2A^−/−^ and E2A^−/−^HEB^−/−^ knockout ES cells stained for pluripotency markers in LIF/serum (L/S). **(B)** qRT-PCR analysis of parental and knockout cell lines in L/S and differentiated in N2B27 over 5 days. Error bars represent standard deviation of a minimum of 3 biological replicates. *p<0.05 by Student’s t-test. **(C)** Cells differentiated in N2B27 for 5 days stained for Oct4 and Sox1, and **(D)** the mature neuronal marker, Tuj1, and the mesendodermal marker, T-brachyury (T-bra).

We next examined the ability of the knockout cells to differentiate under standard neural monolayer conditions. qRT-PCR analyses showed that, compared to controls, E2A^−/−^ and E2A^−/−^HEB^−/−^ cells were unable to robustly upregulate markers of neural differentiation, including *Sox1, Foxd4* and *Pax3* (***Figure 3B and Supplementary Figure 5***). Furthermore, whilst knockout cells were able to downregulate expression of the naïve pluripotency marker, *Nanog*, they maintained comparably high expression of the epiblast markers *Oct4* and *Fgf5*. Immunostaining of differentiation cultures for Sox1, Oct4 (***Figure 3C***) and the mature neuronal marker Tuj1 (***Figure 3D***) further supported the gene expression data. Interestingly, we also observed that disruption of E-protein expression resulted in an upregulation of the mesendodermal lineage markers, *T-brachyury* and *Eomes* (***Figure 3B,D***), as well as re-expression of *Nanog*, suggesting that these cells were beginning to adopt an identity more similar to proximal epiblast, despite being cultured under neural differentiation conditions. Furthermore, we noted that both the neural differentiation defect and the upregulation of mesendodermal markers was more pronounced when HEB was deleted in addition to E2A, indicating that HEB is able to functionally compensate for E2A, to a limited extent, in this context. Taken together with the overexpression data, these findings suggest that E2A is both sufficient and required for efficient neural differentiation of ES cells.

### Identification of early E2A response genes

Having identified a novel role for E2A homodimers in driving neural differentiation of ES cells, we next sought to identify downstream targets of E2A in this process. We have previously reported that BMP blocks neural differentiation by maintaining E-cadherin (Malaguti et al., 2013), and others have reported that E2A is a direct transcriptional repressor of E-cadherin (Pérez-Moreno et al., 2001). We therefore hypothesised that E2A homodimers may also repress E-cadherin protein expression to enable neural differentiation. However, contrary to this hypothesis, we detected no significant downregulation of E-cadherin expression in response to induction of E2A homodimers (***Supplementary Figure 6A***), and therefore conclude that E-cadherin is unlikely to be a direct target of E2A in this context.

To identify novel downstream targets and examine genome-wide changes in expression in response to induction of E2A homodimers, an RNA-sequencing (RNA-seq) approach was chosen. One aim was to identify factors that mediate the ability of E2A to upregulate Sox1. Expression of *Sox1* transcript was robustly upregulated by 18h compared to the corresponding no dox control, whilst expression of Sox1 protein was first detected at 24h (***Supplementary Figure 6B,C***). Based on these observations we performed RNA-seq analysis at 0h (no dox), 18h and 24h timepoints in triplicate to capture changes in genes acting upstream of Sox1. We found that of the 335 genes that were upregulated at 18h compared to 0h, 291 remained significantly upregulated at 24h, whilst 112 out of the 174 genes downregulated at 18h remained downregulated at 24h (***Figure 4A***).

**Figure 4.**
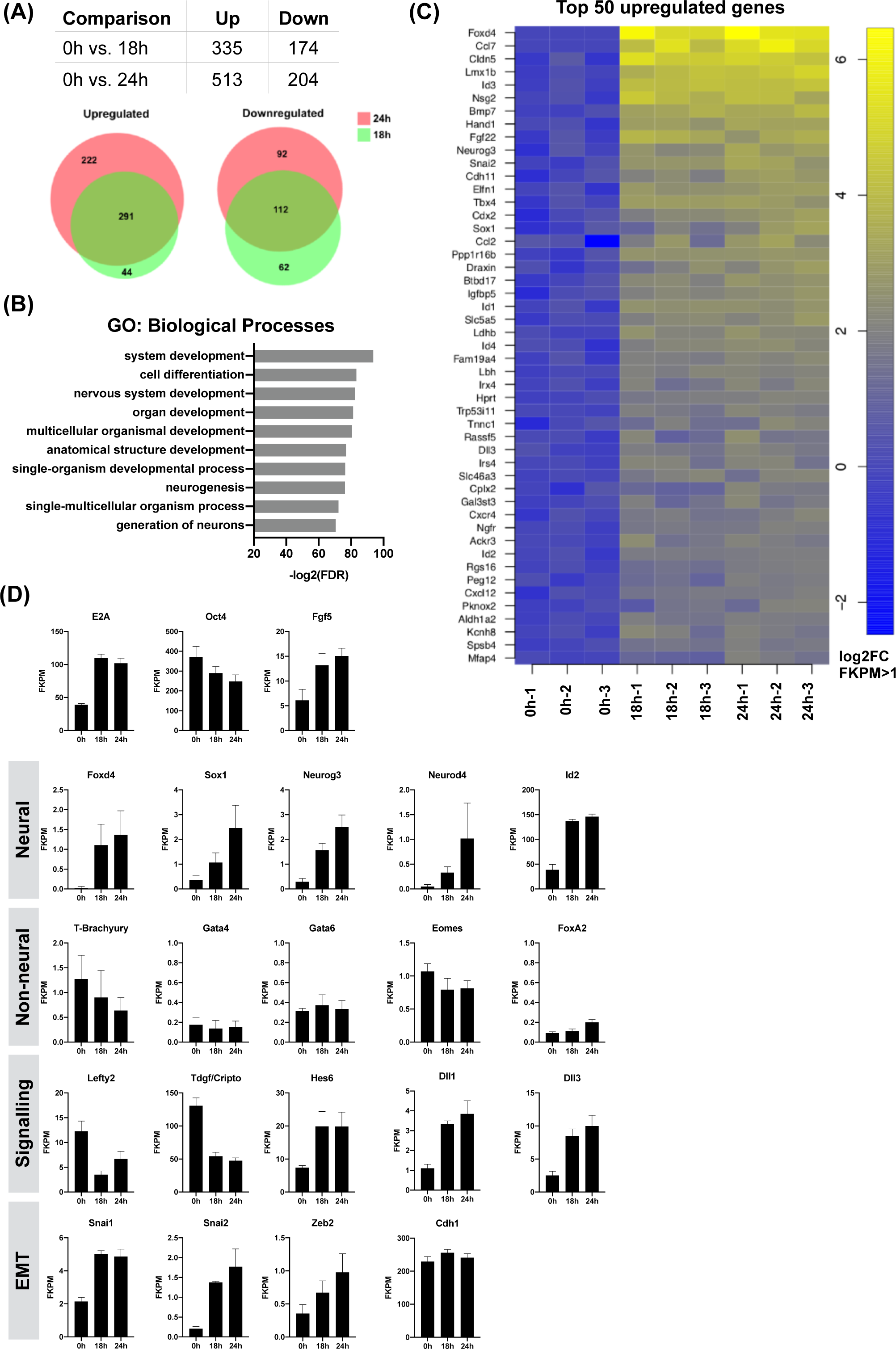
Global transcriptome profiling by RNA-seq reveals E2A homodimers specifically upregulate neural lineage markers. Differentially expressed genes in each comparison according to threshold on minimum log2 fold-change (2) and maximum false discovery rate (0.05). Proportional Venn diagrams illustrate overlap of differentially expressed genes between timepoints. **(B)** Gene Ontology analysis of top 50 upregulated genes at 24h. **(C)** Heatmap of top 50 genes upregulated at 24h. **(D)** Mean FKPM RNA-seq expression values for selected genes at 0h, 18h and 24h. Error bars represent standard deviation of 3 biological replicates.

### E2A homodimers specifically promote neural lineage commitment

Gene ontology (GO) analysis of genes upregulated at 24h revealed an enrichment for terms associated with neural differentiation, including ‘nervous system development’, ‘neurogenesis’ and ‘generation of neurons’ (***Figure 4B***). Further interrogation of the RNA-seq data highlighted an upregulation of neural lineage-associated genes, including *Foxd4* (log2 fold-change (logFC) 5.7; false discovery rate (FDR) 0.0003), *Sox1* (logFC 2.8; FDR 0.003), *Neurog3* (logFC 3.1; FDR 7.18E-05) and *Neurod4* (logFC 4.4; FDR(0.02 (***Figure 4C,D***). In contrast, early markers of non-neural lineages such as *T-brachyury, Eomes, Gata4, Gata6* and *FoxA2* were not upregulated, suggesting that E2A homodimers specifically drive neural lineage commitment in ES cells. We additionally observed an upregulation of EMT-related genes, including *Snai1, Snai2* and *Zeb2*, but no detectable early downregulation of *E-cadherin*, in line with our previous qRT-PCR analyses. We also found an upregulation of Notch signalling components including *Dll1* and *Dll3*, the FGF target gene *Hes6* (Koyano-Nakagawa et al., 2000; Murai et al., 2007), and downregulation of Nodal pathway genes, including the Nodal co-receptor, *Cripto* (*Tdgf1*), and the Nodal target gene, *Lefty2*.

### E2A drives neural commitment by upregulating Foxd4 and dampening Nodal signalling

The forkhead box transcription factor, Foxd4, is a likely candidate for mediating the effect of E2A on neural differentiation. Foxd4 was identified by RNA-seq as the most highly upregulated gene at 24 hours following E2A homodimer induction (***Figure 4C***), and is required for neural fate acquisition in mouse ES cells (Sherman et al., 2017). This is consistent with our observation that *Foxd4* is transiently upregulated at day 3 of neural differentiation, prior to overt *Sox1* expression, whilst E2A^−/−^ and E2A^−/−^ HEB^−/−^ cells fail to upregulate *Foxd4 (****Figure 3B***).

Alternatively, E2A could be modulating differentiation by dampening Nodal activity. The Nodal pathway genes *Cripto* and *Lefty2* were found to be downregulated in response to E2A induction (***Figure 4D***). During gastrulation, the anterior epiblast fated for neuroectoderm is ‘silent’ for Nodal signalling (Peng et al., 2016), and inhibition of the pathway has been shown to promote neural differentiation in both mouse and human ES cells (Camus et al., 2006; Vallier et al., 2004; Watanabe et al., 2005). We found that whilst both *Cripto* and *Lefty2* were rapidly downregulated and maintained at basal levels during differentiation of control cells, they were progressively upregulated in E-protein knockout lines, with a more dramatic effect evident in E2A^−/−^HEB^−/−^ cells (***Figure 5A***).

**Figure 5.**
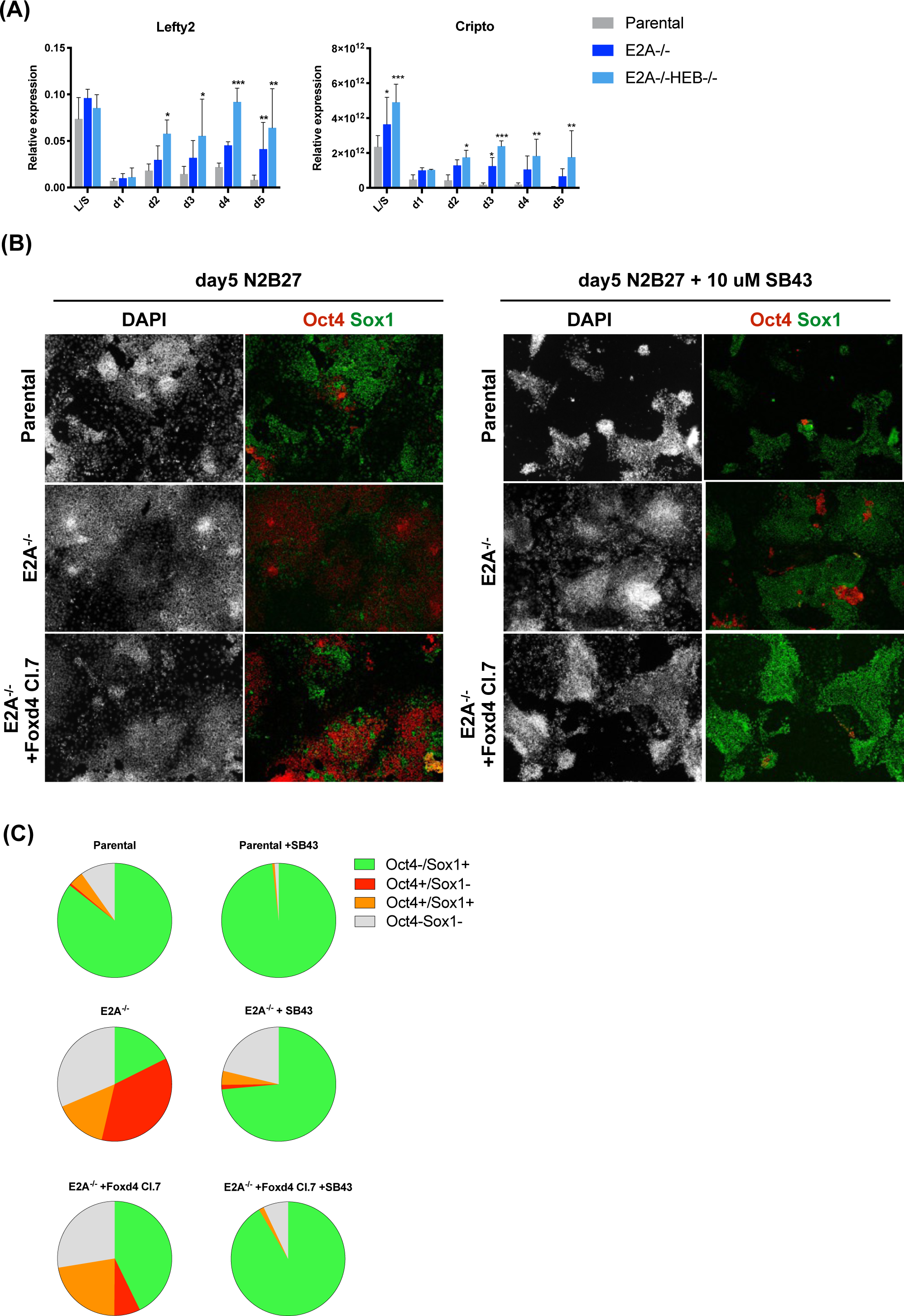
Combined Foxd4 expression and Nodal signalling inhibition rescues differentiation defect in E-protein knockout cells. **(A)** qRT-PCR analysis of Nodal pathway gene expression in knockout and parental ES cell line differentiation. Error bars represent standard deviation of a minimum of 3 biological replicates. *p<0.05 by Student’s t-test. **(B)** Immunostaining of parental and knockout cells at day 5 of differentiation +/-10 uM SB43. **(C)** Quantification of immunostaining of cells at day 5 of differentiation +/-10 uM SB43 performed by quantitative image analysis. Shown are the mean values for 3 biological replicates. A minimum of 8000 nuclei were scored per experiment. Data for all replicates are shown in Supplementary Table 3.

Based on these observations we hypothesised that Foxd4 and/or a dampening of Nodal signalling could explain how E2A can drive neural differentiation. To test this, we sought to determine whether ectopic expression of Foxd4 or inhibition of Nodal signalling could rescue the neural differentiation defect observed in E-protein knockout cells.

We first generated E2A^*−/−*^ ES cell lines that ectopically express Foxd4. A plasmid encoding Foxd4 cloned upstream of an IRES-puromycin resistance cassette, under the control of a PGK promoter, was assembled and transfected into E2A^*−/−*^ cells. *Foxd4* expression was assessed in undifferentiated ES cells by qRT-PCR (***Supplementary Figure 7A***) and three clonal lines were chosen for subsequent analyses (clones 3, 7 and 8). Ectopic expression of Foxd4 was not sufficient to force neural differentiation of E2A^*−/−*^ ES cells cultured in LIF/serum, as assessed by *Sox1* expression (***Supplementary Figure 7A***). Using quantitative immunostaining analysis, we observed that when cells were placed under differentiation conditions, however, ectopic expression of Foxd4 was able to partially restore *Sox1* expression in knockout cells (***Figure 5B,C, Supplementary Figure 7B and Supplementary table 3***), and promote a large proliferation of Oct4-expressing cells present in the culture (***Figure 5B,C***).

We then sought to test whether inhibition of the Nodal pathway using SB431542 (SB43), which inhibits the activity of TGF-β receptors, could restore neural differentiation in E-protein knockout cells. We observed that SB43 treatment increased the proportion of Sox1 expression in control cells, and resulted in a strong upregulation of Sox1 in E2A^−/−^ and E2A^−/−^HEB^−/−^ lines. When applied to cells also ectopically expressing Foxd4, SB43 was able to effect a robust differentiation response (***Figure 5B,C and Supplementary Figure 7B,C***). Taken together, these results suggest an additive effect of ectopic Foxd4 expression and Nodal pathway inhibition in restoring the differentiation capacity of E2A^−/−^ cells. We conclude that E2A homodimers promote neural differentiation by upregulating Foxd4 whilst dampening Nodal activity.

## Discussion

The key extracellular signalling pathways that inhibit differentiation of ES cells are now well established (Haegele et al., 2003; Smith et al., 2008; Ying et al., 2003a; Zhang et al., 2010). A considerable amount of progress has also been made in elucidating the network of transcription factors that control later aspects of mammalian neuronal specification and differentiation (Imayoshi and Kageyama, 2014). Much less is known, however, about the molecular processes that link these two events at the earliest stages of neural differentiation. Recent work has identified roles for transcription factors such as Zfp521, Oct6 and Zic1/2 as intrinsic regulators of early neural differentiation (Kamiya et al., 2011; Sankar et al., 2016; Zhu et al., 2014). However, whilst it has been demonstrated that Oct6 is involved in the activation of several neural fate-promoting genes (Zhu et al., 2014), Zfp521 and Zic1/2 are more likely playing a role in consolidating, rather than initiating, neural fate (Iwafuchi-Doi et al., 2012; Kamiya et al., 2011). In this study, we sought to identify novel intrinsic regulators of neural fate commitment in ES cells, and to elucidate the molecular events unfolding downstream of the BMP signalling pathway during this process.

The HLH transcription factor Id1 has previously been identified as a key effector of the BMP pathway, and its overexpression has been shown to block entry into the neural lineage and promote differentiation to alternative fates (Malaguti et al., 2013; Ying et al., 2003). Whilst the function of Id proteins as dominant negative inhibitors of bHLH transcription factors is well characterised in heterologous systems (Norton et al., 1998), it has not previously been investigated whether Id1 could also be functioning via this classical mechanism during neural commitment. Taken together with the observation that E2A is the major binding partner of Id1 in ES cells (Davies et al., 2013), we therefore hypothesised that Id1 maintains pluripotency by sequestering E2A, preventing it from forming functional homo- or heterodimers, which could otherwise initiate a differentiation response.

Despite their widespread expression during embryogenesis (Pérez-Moreno et al., 2001), a definitive role for Class I bHLH transcription factors in early neural development has not been explored, with E-proteins often relegated to simply being obligate dimerisation partners for tissue-specific Class II bHLH factors. Single E-protein knockouts in mice have also failed to uncover any robust neurodevelopmental deficiencies, whilst attempts to generate E-protein compound knockouts to overcome the expected problem of functional compensation between family members have thus far been unsuccessful as mice with knockout of even single E-proteins do not survive beyond two weeks after birth (Ravanpay and Olson, 2008). In *Drosophila*, however, loss of the only Class I bHLH factor gene, *daughterless*, has been found to result in notable neural differentiation defects (Caudy et al., 1988).

In this study we explored a role for E2A in neural fate commitment. Generation of an endogenously-tagged E2A-V5 ES cell line enabled us to define the expression of E2A from pluripotency to the onset of differentiation on the single cell level. Using a doxycycline-inducible system, we report that overexpression of E2A is sufficient to promote commitment to the neural lineage in ES cells, even under non-permissive culture conditions. Specifically, we found that forced E2A homodimers, rather than E2A-bHLH heterodimers, appear to drive this process, as introduction of a single amino acid change into E2A to disrupt homodimer stability, whilst still maintaining heterodimerisation capacity (Benezra, 1994), did ablate the ability of the forced homodimer to promote *Sox1* expression. This was particularly striking as a physiological role for E2A homodimers has not previously been described outside of the context of B-cell specification (Shen and Kadesch, 1995). Interestingly, we also found that overexpression of E2A was able to drive upregulation of its own negative regulator, *Id1*, consistent with previous descriptions of a Class I/Class V HLH feedback loop in *Drosophila*, where overexpression of the E-protein *daughterless* causes transcriptional upregulation of the Id protein, *extramacrochaetae* (Bhattacharya and Baker, 2011).

The findings from the gain-of-function assays were further supported by data from the loss-of-function studies. Generation of novel E2A^−/−^ and E2A^−/−^HEB^−/−^ ES cell lines highlighted that whilst knockout cells were able to self-renew normally, they were not able to differentiate efficiently. Additionally, although knockout cells were able to downregulate *Nanog*, they maintained relatively high expression of epiblast markers *Fgf5* and *Oct4*. This suggests that whilst knockout cells are able to navigate the exit from pluripotency efficiently, they fail to complete the proposed second stage of neural lineage commitment involving the transition from epiblast-like cells to neuroectoderm-like cells (Zhang et al., 2010). The observation that E2A^−/−^HEB^−/−^cells displayed a more pronounced neural differentiation defect suggests that HEB is, indeed, able to at least partially compensate for loss of E2A in this context. This is in line with previous studies, which have demonstrated that HEB is able to functionally replace E2A during B-cell commitment (Zhuang et al., 1998), highlighting the extent of redundancy between E-protein family members. That E2A^−/−^HEB^−/−^ cells also appear to preferentially upregulate markers of mesendodermal lineages, including *T-brachyury* and *Eomes*, is a potentially interesting topic of investigation for future studies, especially as HEB (and E2A) have previously been implicated in promoting mesendodermal lineage specification, both in *Xenopus* and in human ES cells (Li et al., 2017; Yoon et al., 2011).

In this work we identified Foxd4 as a key gene strongly upregulated in response to forced E2A homodimers, positioning it upstream of Sox1 expression during neural commitment. These data are in line with recent observations also made in ES cells (Sherman et al., 2017), and with the reported expression of *Foxd4* in the mouse neuroectoderm at E7.5 (Kaestner et al., 1995). The *Xenopus* homologue of Foxd4, Foxd4l1 (previously Foxd5), has a well-established role as part of a broader network of neural fate stabilising factors (Yan and Moody, 2009), and it has been shown to expand the population of progenitor cells in the immature neuroectoderm, whilst repressing the transcription of genes associated with neural differentiation (Sullivan et al., 2001). A potential role for Foxd4 in maintaining neuroectodermal cells in a proliferative, non-differentiating state, therefore, may also explain the large proliferation of Oct4-positive cells we observed when Foxd4 was ectopically expressed in E2A^−/−^ cells.

We also reported that components of the Nodal signalling pathway were downregulated upon overexpression of E2A homodimers, and conversely upregulated during differentiation of E-protein knockout cells. Interestingly, E2A has also been shown to repress transcription of *lefty* in *Xenopus*, with knockout of E2A causing upregulation of *lefty* and a subsequent failure in mesendodermal fate specification (Wills and Baker, 2015). We propose that E2A could be playing a similar role to repress Nodal signalling in ES cells, but during neural fate commitment – a process in which inhibition of Nodal signalling is already known to be important (Vallier et al., 2004; Watanabe et al., 2005). We also report that defective E2A^−/−^ neural differentiation can be restored by the combined effect of ectopic Foxd4 expression and Nodal inhibition, suggesting a possible dual role for E2A in this process. In summary, we propose that E2A plays an instructive, rather than passive, role in promoting neural fate commitment in pluripotent cells by promoting the transcription of neural lineage genes whilst simultaneously suppressing the Nodal signalling pathway.

## Acknowledgements

We thank Steve Pollard and Pooran Dewari for help with designing the CRISPR/Cas9 targeting experiments, Thomas Maynard for Foxd4 plasmids, and Edinburgh Genomics for performing library preparation, RNA-sequencing and bioinformatics analysis.

## Methods

### Mouse ES cell culture and neural differentiation

ES cells were maintained in GMEM supplemented with 2-mercaptoethanol, non-essential amino acids, glutamine, pyruvate, 10% foetal calf serum (FCS) and 100 units/mL LIF on gelatinised tissue culture flasks (Smith, 1991). Monolayer neural differentiation was performed as described in (Pollard et al., 2006). Briefly, ES cells were washed in DMEM/F12 to remove all traces of serum and plated at 1×10^4^ cells/cm^2^ in N2B27 medium on gelatinised tissue culture plates. N2B27 consists of a 1:1 ratio of DMEM/F12 and Neurobasal media supplemented with 0.5% N2, 0.5% B27, L-glutamine and 2-mercaptoethanol. For Nodal signalling inhibition experiments, cells were seeded in N2B27 supplemented with 10 µM SB431542.

### CRISPR/Cas9 epitope tagging of ES cell lines

Guide RNA was manually designed to introduce the V5 tag into the 3’UTR of E2A based on annotation of the final coding exon and the 3’UTR sequence. A ~200bp sequence around the stop codon was used as an input for guide RNA design using either crispr.mit.edu or crispor.tefor.net web-based design tools. High-scoring guide RNAs were chosen based on minimal predicted off-target cleavage events, and having a cut site within the 3’UTR, preferably within 10bp of the stop codon. For ssODN design, the V5 tag sequence was flanked by ~75bp homology arms and a PAM-blocking mutation (NGG>NGC) was introduced into the 3’UTR sequence. Ribonucleoprotein (RNP) complexes were assembled as described in (Dewari et al., 2018). Briefly, crRNA and tracrRNA oligos were mixed in equimolar concentration, heated at 95°C for 5 min and allowed to cool to room temperature to anneal. 10 µg of recombinant Cas9 protein was added to the annealed cr/tracrRNAs to form the RNP complex, which was incubated at room temperature for 20 min and stored on ice until electroporation. 30 picomoles of ssODN was added to the RNP complex immediately before electroporation. 5×10^4^ ES cells were transfected using the 4D Amaxa nucleofection system (Lonza) using the optimised programme, CA-210. Following transfection, cells were transferred into a 6-well plate and allowed to recover for 3-5 days in ES cell media. Clonal cell lines were isolated from bulk populations by manual colony picking and were subsequently analysed for successful knock-in by PCR genotyping, immunocytochemistry and Sanger sequencing of a ~500bp region spanning the target site.

Synthetic Alt-R crRNA, tracrRNA, ssODN Ultramer template DNA oligonucleotides and recombinant Cas9 protein were manufactured by Integrated DNA Technologies (IDT), USA. Guide RNA and ssODN DNA sequences are provided in ***Supplementary table 1***.

### Knockout ES cell line generation

For generation of E2A^−/−^ ES cells, guide RNAs were designed to target exon 3 of the E2A gene. Targeting was performed on the E2A-V5 cell line to facilitate knockout line validation by loss of V5 signal. For generation of E2A^−/−^HEB^−/−^ double knockout ES cells, exon 9 of the *HEB* locus was targeted in E2A^−/−^ knockout cells (clone 9) in order to disrupt both HEBcan and HEBalt splice variants. Guides were designed, assembled as RNPs with recombinant Cas9, and transfected as detailed above. Knockouts were verified by Sanger sequencing of targeted allele and western blot analysis using V5 (Thermo Scientific), and HEB (Abcam) antibodies. Experiments presented in main text were performed using E2A^−/−^ clone 9 and E2A^−/−^HEB^−/−^ clone 12.

### Doxycycline-inducible E2A cell lines

The E47 monomer construct is comprised of an N-terminally FLAG-tagged mouse E47 sequence. For generation of the E47 forced homodimer construct the DNA sequence corresponding to the FLAG-tagged mouse E47 was tethered at its C-terminus to the N-terminus of a second mouse E47 sequence through a 13-amino acid flexible linker sequence of TGSTGSKTGSTGS by overlapping extension PCR. For generation of the C570A homodimerisation-deficient cell lines, mutations were introduced into both E47 sequences in the forced dimer construct using PCR site-directed mutagenesis and verified by Sanger sequencing. Inducible cells lines were generated using a inducible cassette exchange (ICE) technique (Iacovino et al., 2013). Cells were stimulated with 1 *µ*g/mL doxycycline in all experiments.

### Generation of Foxd4 rescue lines

The full-length Foxd4 cDNA sequence was cloned into a pPGK-IRES-puromycin resistance plasmid. E2A^−/−^ (clone 9) cells were transfected with the resulting construct using the 4D Amaxa nucleofection device and clonal lines were isolated by puromycin selection (1 µg/mL). Following clonal line expansion, ES cells were characterised by qRT-PCR for *Foxd4* expression.

### Immunohistochemistry, western blot analysis and flow cytometry

Samples were fixed with 4% paraformaldehyde for 10 minutes at room temperature and incubated with blocking buffer (PBS, 0.1% Triton X-100, 3% donkey serum) for 30 minutes at room temperature. Primary antibodies were diluted in blocking buffer and incubated with cells overnight at 4°C. After three washes with PBS, cells were incubated with secondary antibodies conjugated with Alexa fluorophores (Life Technologies) diluted 1:1000 in blocking buffer for 1 hour at room temperature. For nuclear counterstaining cells were incubated with 1 µg/mL DAPI (Sigma) for 5 minutes following immunostaining. Cells were washed a minimum of three times in PBS and imaged on an Olympus inverted fluorescence microscope. Images were analysed in Fiji.

Primary antibodies used for immunocytochemistry were:- β-III-tubulin/Tuj1 (1:1000; Biolegend), Nanog (1:200 eBioscience), Oct4 (1:200; Santa Cruz), Sox1(1:200; BD Pharmigen), Sox2 (1:200; Abcam) and V5 (1:200-500; Thermo Scientific).

Primary antibodies used in western blot analysis were E2A (1:200; sc-416, Santa Cruz), HEB (1:200; sc-28364), V5 (1:500; Thermo Scientific) and β-III-tubulin/Tuj1 (1:1000; Biolegend). Blots were probed using horseradish peroxidase-conjugated secondary antibodies.

For flow cytometry analysis cells were stained with E-cadherin (CD324) DECMA-1 (1:300; eBioscience). Flow cytometry was performed on the BD Accuri C6 and analysis was performed using FlowJo software.

### Quantification of immunostaining

Immunofluorescence was quantified by nuclear segmentation based on DAPI signal and manual editing of segmentation results using NesSys software, as detailed in (Blin et al., 2019).

### qRT-PCR

Total RNA was isolated from cells using the Absolutely RNA Miniprep Kit (Agilent). cDNA was generated using Maloney murine leukaemia virus reverse transcriptase (MMLV-RT) and random primers. Primers and UPL probes (Roche) used are detailed in ***Supplementary table 2***. All gene expression values are normalised to the housekeeping gene *SDHA*.

### RNA sequencing

RNA was isolated from cells as previously described and RNA quality was verified using an Agilent 2100 Bioanalyzer. Subsequent cDNA library preparation, sequencing and bioinformatics analysis, including differential gene expression analyses, were performed by the Edinburgh Genomics facility. Library preparation was performed using the TruSeq Stranded mRNA Library Prep Kit (Illumina) and libraries were sequenced on the Illumina HiSeq4000. Reads were mapped to the mouse genome GRCm38 from Ensembl and were aligned to the reference genome using STAR (version 2.5.2b). Differential gene analysis was carried out with edgeR (version 3.16.5). Differentially expressed genes were assigned based on a minimum fold-change of 2 and a maximum false discovery rate (FDR) of 0.05 Gene ontology analysis was performed on genes upregulated at 24h using the STRING database (Szklarczyk et al., 2017). RNA-seq data will be deposited in the GEO database.

### Statistical analyses

qRT-PCR data are represented as mean ±SD for a minimum of three experimental replicates. Statistical significance was calculated using Student’s *t*-tests for pairwise comparisons, or ANOVA with correction for multiple comparisons for two or more samples. Statistically significant differences are shown as follows: *p<0.05, **p<0.01 and ***p<0.001.

---

**Supplementary Figure 1.**
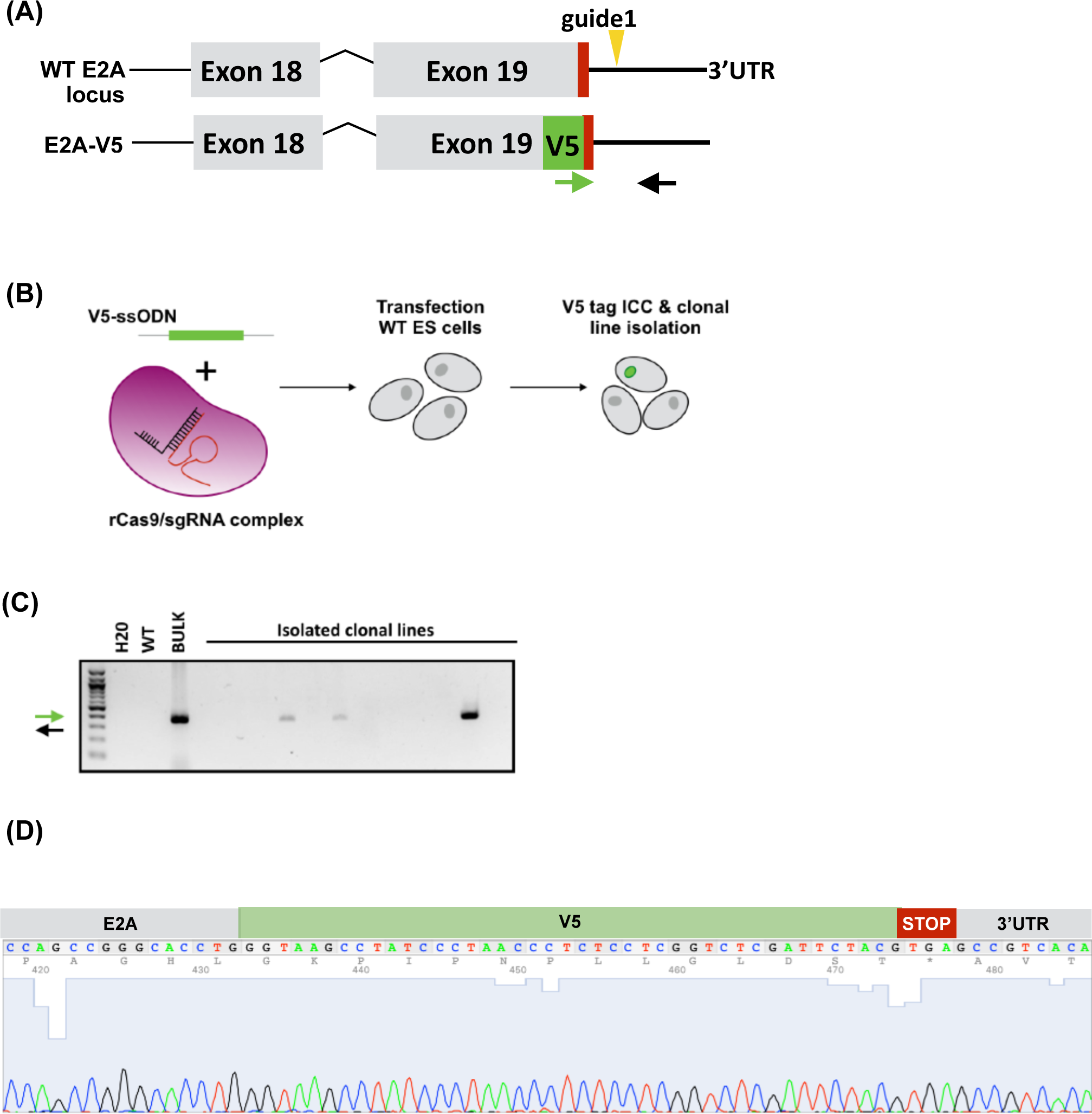
Generation and validation of endogenously tagged E2A-V5 ES cell lines. **(A)** Schematic of exon targeting of E2A to generate tagged cell lines. V5 tag is shown in green, stop codon in red, and yellow arrow indicates location of the guide RNA (sgRNA) sequence. **(B)** Method used to knock in the V5 tag by nucleofection of a complexed ribonucleoprotein (RNP) comprised of recombinant Cas9 protein (rCas9) and guide RNA into wild-type ES cells. **(C)** PCR genotyping of derived clonal lines using primer pairs indicated by green and black arrows depicted in (A). Bands of correct size in clonal lines shown, compared to blank (H20), parental WT line, and bulk transfected population. **(D)** Sanger sequencing trace of correctly targeted E2A-V5 clonal lines, confirming correct in-frame insertion of the V5 tag.

**Supplementary Figure 2.**
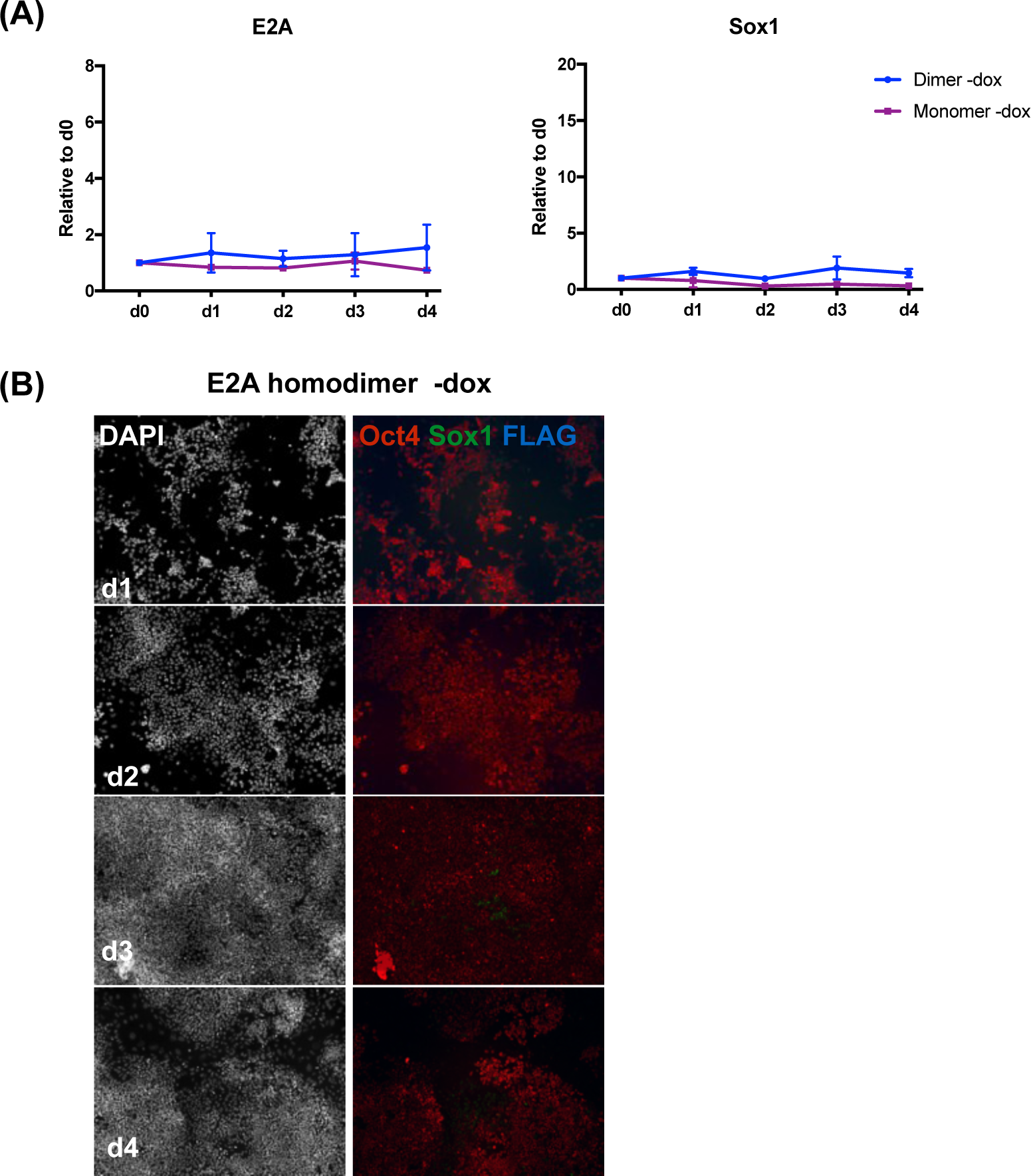
Sox1 is not expressed when inducible cells are cultured in LIF/serum without dox induction. **(A)** qRT-PCR analysis of dox-inducible E2A monomer and forced homodimer cells cultured in LIF/serum without dox. Expression values are normalised to day 0. Error bars represent mean +/-SD of 2 biological replicates. **(B)** Immunocytochemistry of inducible forced homodimer cells cultured in LIF/serum without dox.

**Supplementary Figure 3.**
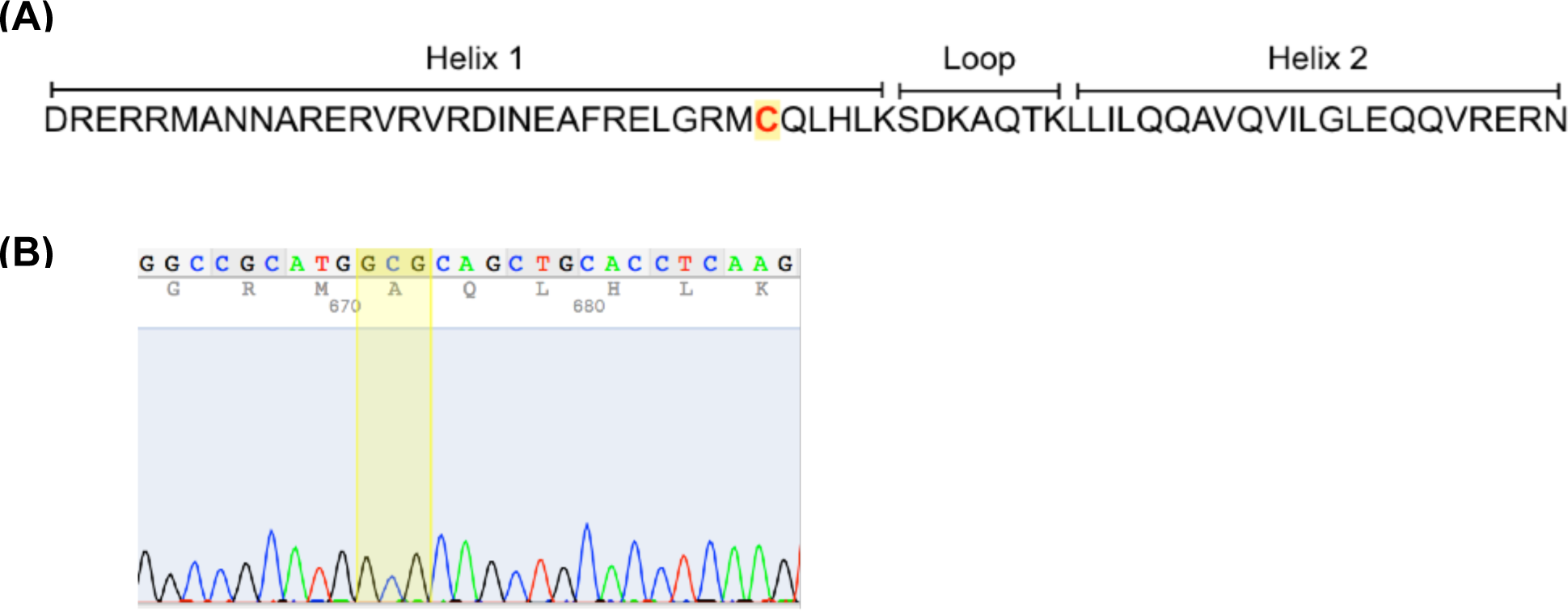
Validation of Cysteine-570 to Alanine (C570A) mutation in forced homodimer cells. **(A)** Sequence of the helix-loop-helix (HLH) domain of wild-type E2A with Cysteine-570 residue in helix 1 highlighted in yellow. **(B)** Sanger sequencing trace of Cys>Ala (C570A) mutation introduced into both E2A monomer sequences in the forced homodimer construct to generate mutant forced homodimer inducible lines (C570A-1 and C570A-9).

**Supplementary Figure 4.**
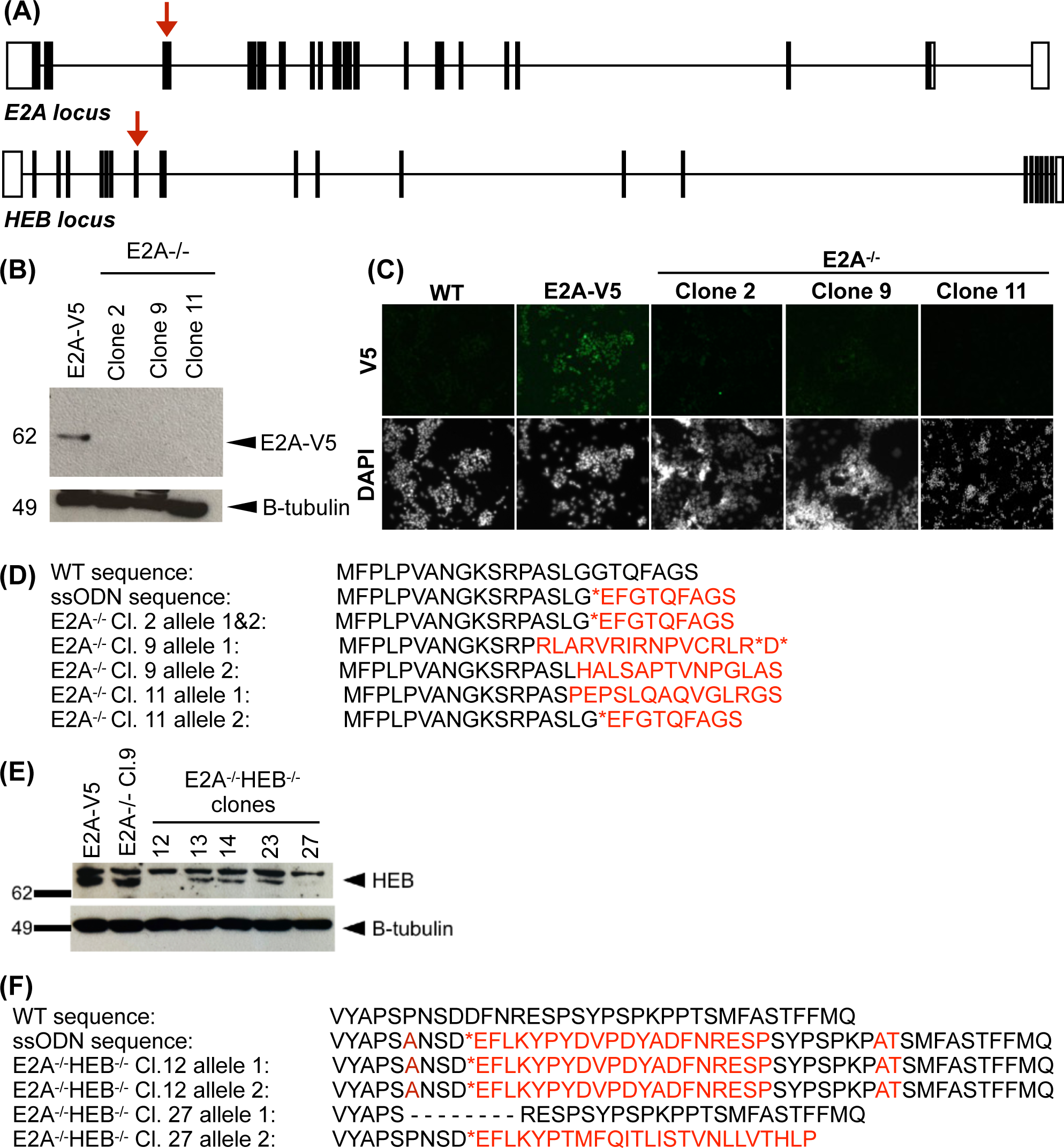
Validation of E-protein knockout lines. (**A**) Structure of *E2A* and *HEB* loci. Red arrow indicates Cas9/sgRNA target site. (**B**) Western blot of E2AV5 Cl.74 (parental cell line) and three E2A knockout clones 2, 9 & 11 using anti-V5 antibody. E2A-V5=69kDa. B-tubulin was analysed as a loading control. (**C**) Immunostaining of WT, E2A-V5 (parental) and E2A−/− clonal lines with anti-V5 antibody. (**D**) Sanger sequencing of E2A−/− clones 2, 9 and 11. WT exon 3 and ssODN template DNA amino acid sequences shown with mutations highlighted in red. E2A−/− clone 2 has biallelic knock-in of the stop codon-containing ssODN template sequence, clone 9 has two independently disrupted E2A alleles and clone 11 has heterozygous knock-in of the ssODN and a second non-homologous end joining (NHEJ)-disrupted allele. (**E**) Western blot of E2AV5 Cl74, E2A−/− clone 9 (parental) and E2A−/−HEB−/− clonal lines using anti-HEB antibody. HEB=85kDa. B-tubulin was analysed as a loading control. (**F**) Sanger sequencing of E2A−/−HEB−/− clones 12 and 27. Targeted WT exon 9 and ssODN template DNA amino acid sequences shown above, with mutations highlighted in red. E2A−/−HEB−/− clone 12 has biallelic knock-in of the ssODN sequence, and clone 27 has two independently disrupted HEB alleles; one allele contains an 8 amino acid deletion, the other allele contains a premature stop codon.

**Supplementary Figure 5.**
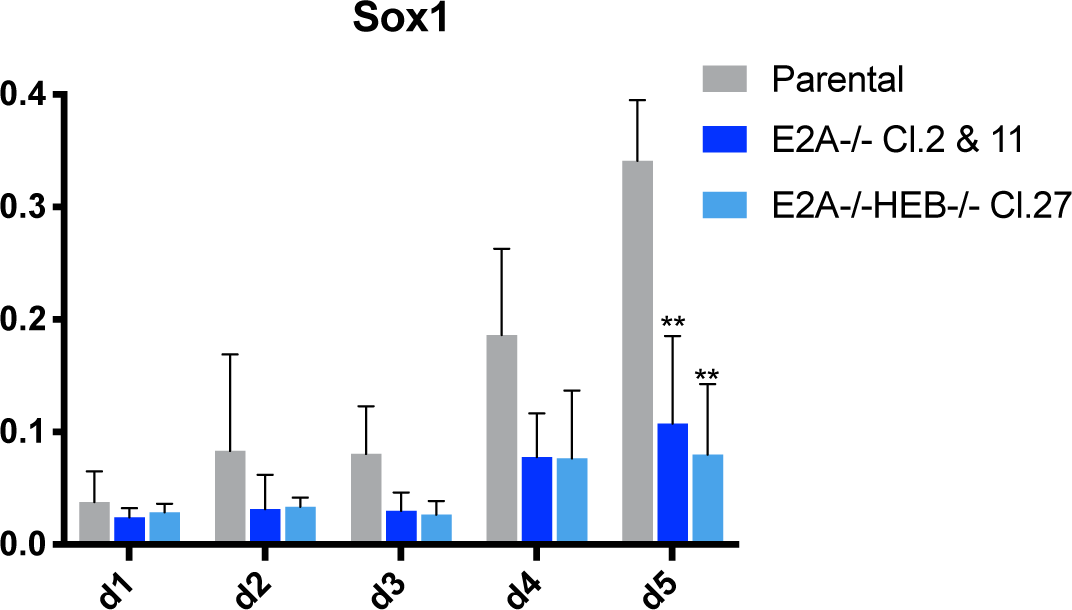
Transcriptional analysis of additional E-protein knockout clones. qRT-PCR analysis of two additional E2A−/− clones (clones 2 and 11) and one additional E2A−/−HEB−/− clone (clone 27) differentiated in N2B27. Relative expression values shown are the mean of three independent experiments for each clonal line.

**Supplementary Figure 6.**
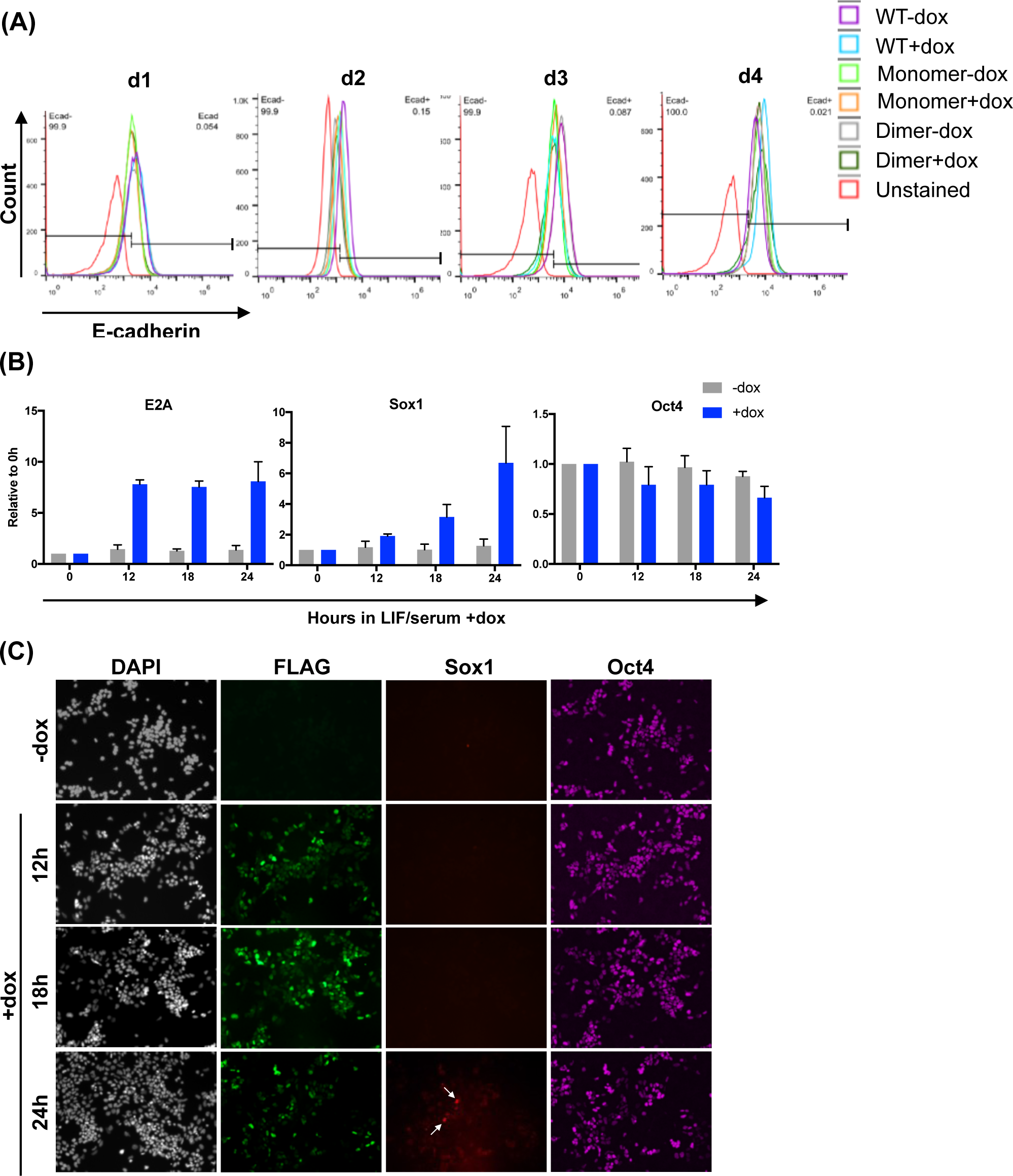
Identifying a time frame for detection of early E2A homodimer response genes. (**A**) Flow cytometric analysis of the effect of overexpression of E2A monomers and homodimers on E-cadherin (E-cad) protein expression. WT ES cells are included as a control and were used in combination with unstained WT cells to define gates. Experiments were repeated twice and the data from replicates follow the same trend as displayed in the figure. (**B**) qRT-PCR analysis of cells following 12, 18 and 24 hours of E2A homodimer induction in LIF/serum. No dox controls are also shown. Error bars represent standard deviation of three biological experiments, which were used for subsequent RNA-sequencing analysis. (**C**) Immunostaining of cells during the 24h timecourse to assess transgene activation using an anti-FLAG antibody, and co-staining with anti-Sox1 and anti-Oct4. White arrows highlight small number of cells that express Sox1 protein 24 hours post-induction.

**Supplementary Figure 7.**
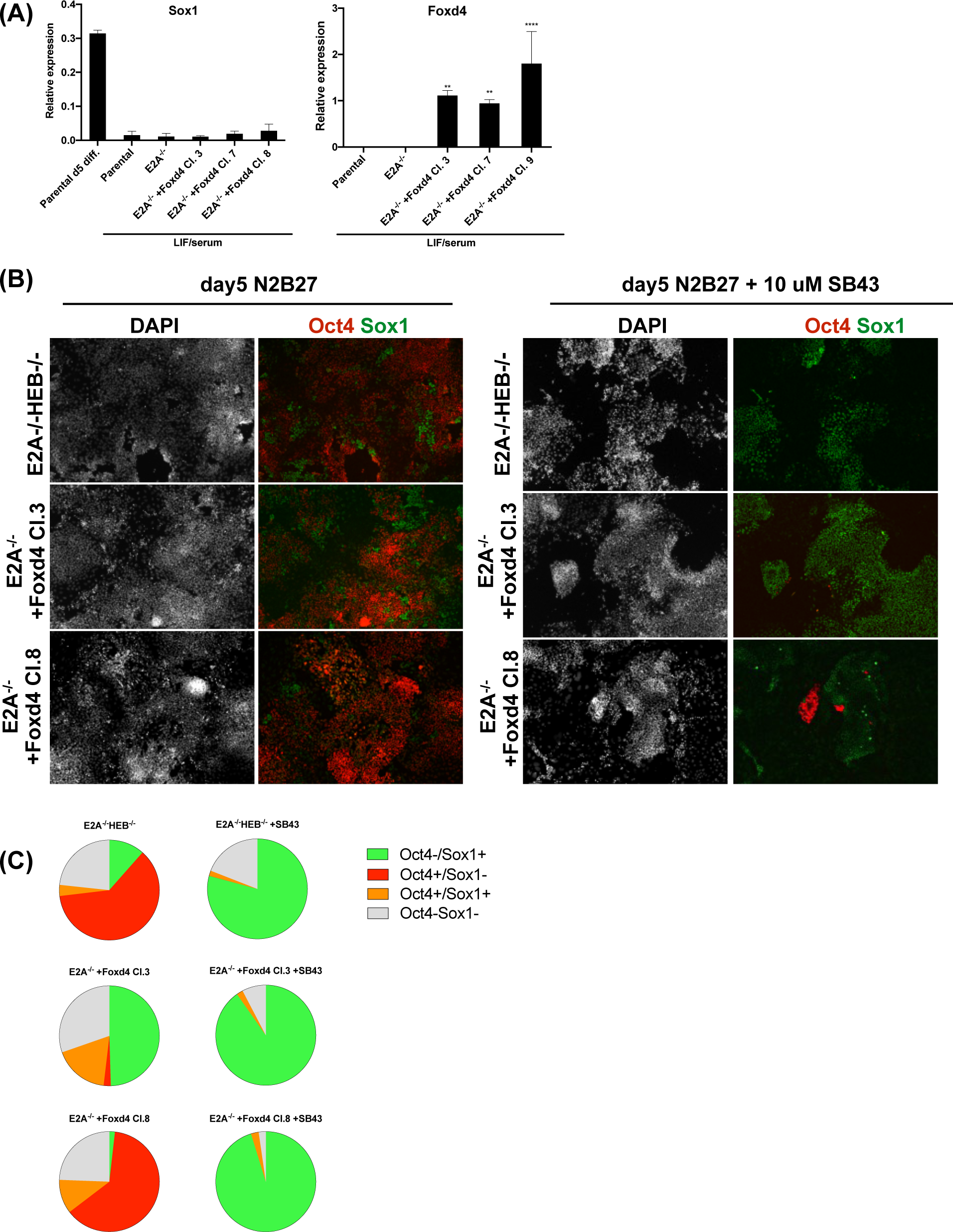
Analysis of additional clonal lines ectopically overexpressing Foxd4 in an E2A−/− background. (**A**) qRT-PCR analysis of knockout and Foxd4 rescue cell lines in LIF/serum. Relative expression values shown are the mean+/-SD of three biological replicates. (**B**) Immunostaining of additional knockout and Foxd4 rescue cell lines differentiated in N2B27+/-SB43. (**C**) Quantification of immunostaining of cells at day 5 of differentiation +/- 10 uM SB43 performed by nuclear segmentation and quantitative image analysis. Shown are the mean values for three independent biological replicates. A minimum of 8000 nuclei were scored per experiment. Data for all replicates are shown in Supplementary table 3.

**Supplementary table 1.**
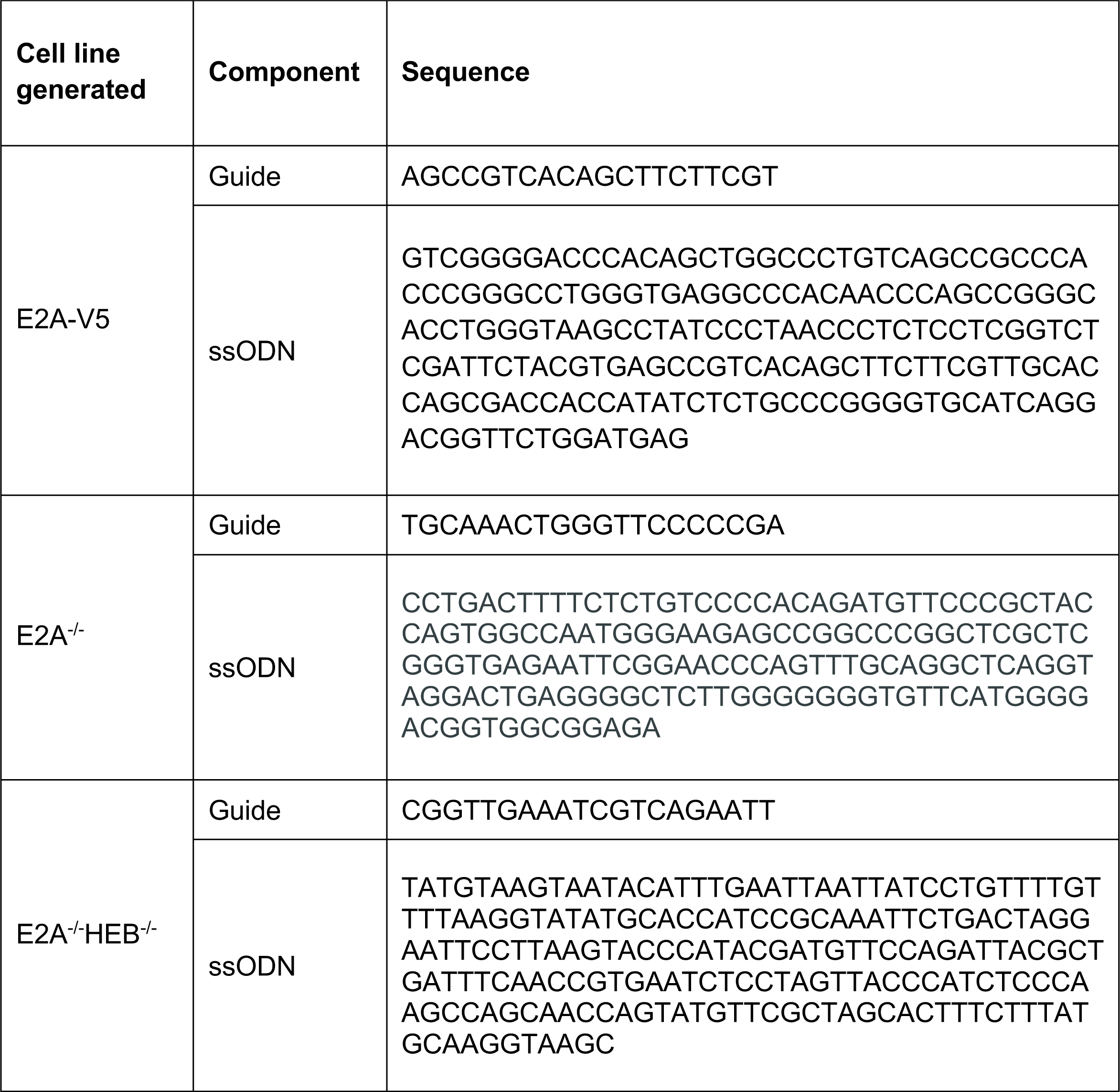
CRISPR/Cas9 targeting guide RNA and ssODN template sequences.

**Supplementary table 2.**
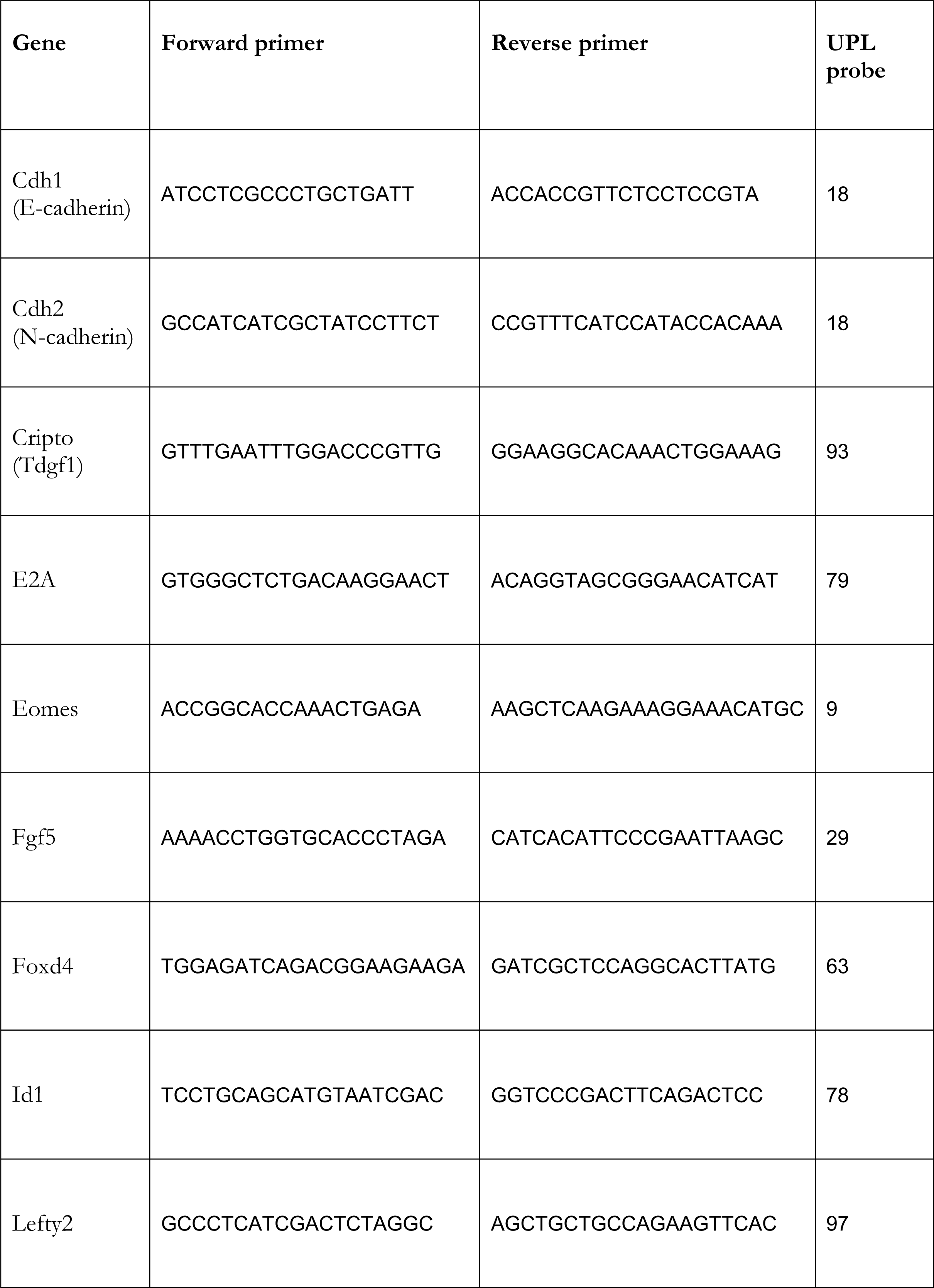
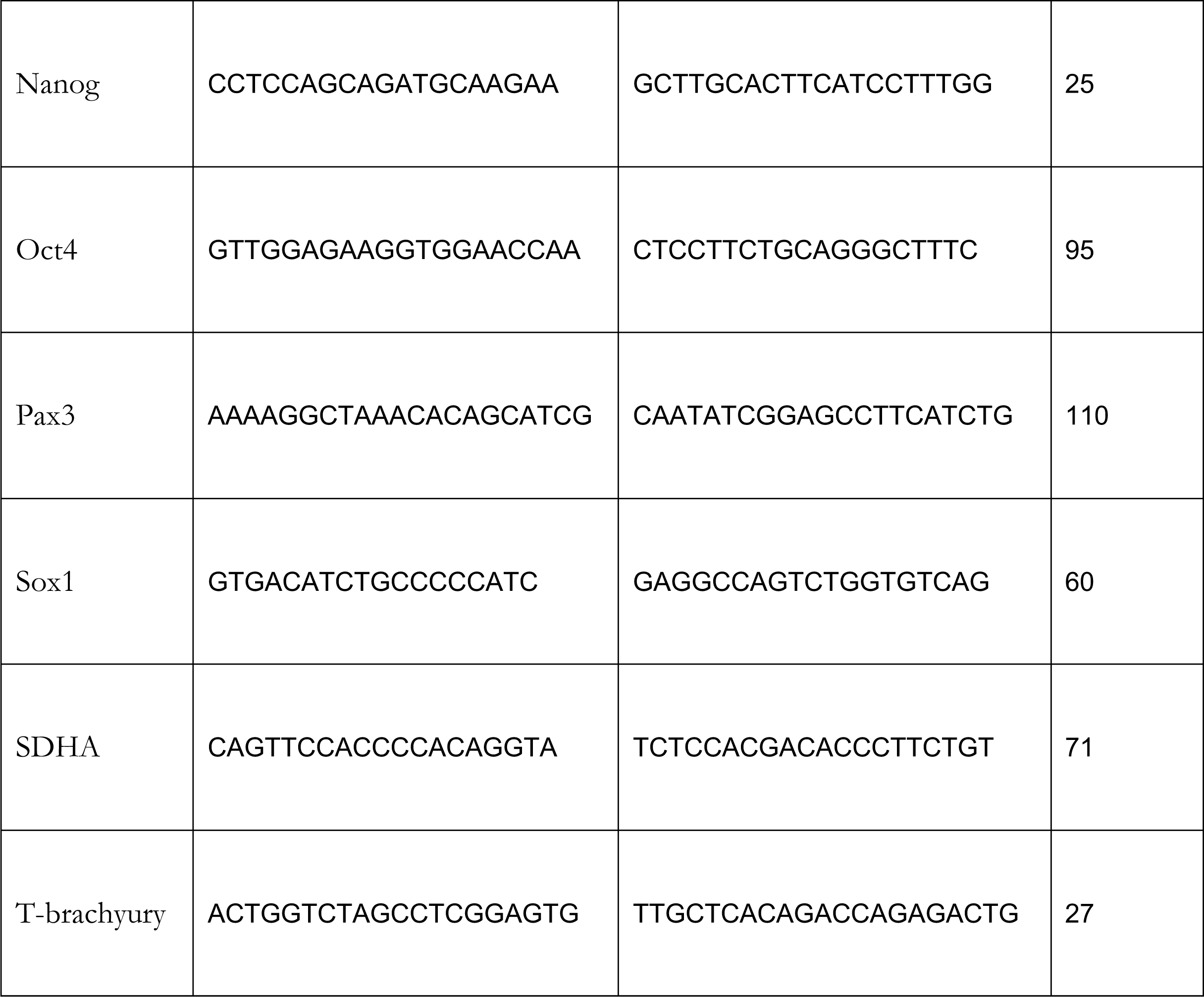
qRT-PCR primers and probes.

**Supplementary table 3.** Immunostaining quantification. (associated with Figure 5) **(A)** Cells differentiated in N2B27 **(B)** Cells differentiated in N2B27+SB43

| **-SB43** | **% Rep1** | **% Rep2** | **% Rep3** | **Mean %** | **Standard deviation** |
| --- | --- | --- | --- | --- | --- |
| **Parental** |  |  |  |  |  |
| **Oct4-/Sox1+** | 72.9 | 89.2 | 83.8 | 82.0 | 8.34 |
| **Oct4+/Sox1-** | 1.8 | 0.0 | 1.5 | 1.1 | 0.98 |
| **Oct4+/Sox1+** | 6.2 | 3.1 | 5.2 | 4.8 | 1.60 |
| **Oct4-Sox1-** | 19.1 | 7.7 | 9.5 | 12.1 | 6.13 |
| **E2A-/-** |  |  |  |  |  |
| **Oct4-/Sox1+** | 4.7 | 7.6 | 56.5 | 22.9 | 29.07 |
| **Oct4+/Sox1-** | 40.3 | 45.6 | 3.9 | 30.0 | 22.68 |
| **Oct4+/Sox1+** | 3.1 | 18.2 | 12.1 | 11.1 | 7.59 |
| **Oct4-Sox1-** | 51.9 | 28.6 | 27.5 | 36.0 | 13.76 |
| **E2A-/-HEB-/-** |  |  |  |  |  |
| **Oct4-/Sox1+** | 17.3 | 6.6 | 56.0 | 26.6 | 28.78 |
| **Oct4+/Sox1-** | 33.4 | 69.6 | 2.8 | 35.3 | 33.95 |
| **Oct4+/Sox1+** | 3.3 | 3.3 | 6.8 | 4.4 | 3.91 |
| **Oct4-Sox1-** | 46.1 | 20.5 | 34.4 | 33.7 | 14.03 |
| **E2A-/- +Foxd4 Cl.3** |  |  |  |  |  |
| **Oct4-/Sox1+** | 57.0 | 61.3 | 43.5 | 53.9 | 9.27 |
| **Oct4+/Sox1-** | 25.5 | 1.0 | 0.9 | 9.1 | 14.19 |
| **Oct4+/Sox1+** | 11.1 | 21.4 | 16.7 | 16.4 | 5.15 |
| **Oct4-Sox1-** | 6.4 | 16.3 | 38.9 | 20.5 | 16.64 |
| **E2A-/- +Foxd4 Cl.7** |  |  |  |  |  |
| **Oct4-/Sox1+** | 42.8 | 45.4 | 40.0 | 42.8 | 2.68 |
| **Oct4+/Sox1-** | 7.1 | 7.7 | 7.0 | 7.3 | 0.38 |
| **Oct4+/Sox1+** | 8.6 | 17.7 | 28.8 | 18.4 | 10.12 |
| **Oct4-Sox1-** | 41.5 | 29.1 | 24.1 | 31.5 | 8.96 |
| **E2A-/- +Foxd4 Cl.8** |  |  |  |  |  |
| **Oct4-/Sox1+** | 10.3 | 0.3 | 1.9 | 4.2 | 5.39 |
| **Oct4+/Sox1-** | 21.5 | 67.6 | 64.8 | 51.3 | 25.83 |
| **Oct4+/Sox1+** | 5.6 | 6.4 | 16.4 | 9.5 | 6.00 |
| **Oct4-Sox1-** | 62.5 | 25.8 | 16.9 | 35.1 | 24.19 |

(B)
| +SB43 | % Rep1 | % Rep2 | % Rep3 | Mean % | Standard deviation |
| --- | --- | --- | --- | --- | --- |
|  | Parental |  |  |  |  |
| Oct4-/Sox1+ | 93.1 | 99.3 | 97.7 | 96.7 | 3.18 |
| Oct4+/Sox1- | 0.0 | 0.0 | 0.0 | 0.0 | 0.00 |
| Oct4+/Sox1+ | 0.0 | 0.7 | 1.1 | 0.6 | 0.56 |
| Oct4-Sox1- | 6.9 | 0.0 | 1.2 | 2.7 | 3.66 |
|  | E2A-/- |  |  |  |  |
| Oct4-/Sox1+ | 73.1 | 65.0 | 93.9 | 77.3 | 14.88 |
| Oct4+/Sox1- | 3.8 | 0.4 | 0.0 | 1.4 | 2.09 |
| Oct4+/Sox1+ | 3.9 | 4.9 | 1.7 | 3.5 | 1.63 |
| Oct4-Sox1- | 19.2 | 29.7 | 4.4 | 17.8 | 12.70 |
|  | E2A-/-HEB-/- |  |  |  |  |
| Oct4-/Sox1+ | 73.5 | 63.5 | 97.8 | 78.2 | 17.19 |
| Oct4+/Sox1- | 0.0 | 0.2 | 0.0 | 0.1 | 0.09 |
| Oct4+/Sox1+ | 1.7 | 1.9 | 1.3 | 1.7 | 1.19 |
| Oct4-Sox1- | 24.7 | 34.4 | 0.9 | 20.0 | 16.92 |
|  | E2A-/- +Foxd4 Cl.3 |  |  |  |  |
| Oct4-/Sox1+ | 82.8 | 85.1 | 96.7 | 88.2 | 7.44 |
| Oct4+/Sox1- | 0.0 | 0.0 | 0.0 | 0.0 | 0.00 |
| Oct4+/Sox1+ | 3.6 | 1.5 | 2.6 | 2.6 | 1.04 |
| Oct4-Sox1- | 13.5 | 13.4 | 0.6 | 9.2 | 7.40 |
|  | E2A-/- +Foxd4 Cl.7 |  |  |  |  |
| Oct4-/Sox1+ | 85.1 | 94.4 | 97.0 | 92.2 | 6.27 |
| Oct4+/Sox1- | 0.0 | 0.0 | 0.0 | 0.0 | 0.00 |
| Oct4+/Sox1+ | 1.8 | 0.6 | 3.0 | 1.8 | 1.16 |
| Oct4-Sox1- | 13.1 | 5.0 | 0.0 | 6.0 | 6.60 |
|  | E2A-/- +Foxd4 Cl.8 |  |  |  |  |
| Oct4-/Sox1+ | 76.3 | 95.6 | 97.0 | 89.6 | 11.58 |
| Oct4+/Sox1- | 0.0 | 0.0 | 0.0 | 0.0 | 0.00 |
| Oct4+/Sox1+ | 0.0 | 2.3 | 3.0 | 1.8 | 1.58 |
| Oct4-Sox1- | 23.7 | 2.1 | 0.0 | 8.6 | 13.14 |

